# ThermoRawFileParser: modular, scalable and cross-platform RAW file conversion

**DOI:** 10.1101/622852

**Authors:** Niels Hulstaert, Timo Sachsenberg, Mathias Walzer, Harald Barsnes, Lennart Martens, Yasset Perez-Riverol

## Abstract

The field of computational proteomics is approaching the big data age, driven both by a continuous growth in the number of samples analysed *per* experiment, as well as by the growing amount of data obtained in each analytical run. In order to process these large amounts of data, it is increasingly necessary to use elastic compute resources such as Linux-based cluster environments and cloud infrastructures. Unfortunately, the vast majority of cross-platform proteomics tools are not able to operate directly on the proprietary formats generated by the diverse mass spectrometers. Here, we presented ThermoRawFileParser, an open-source, crossplatform tool that converts Thermo RAW files into open file formats such as MGF and the HUPO-PSI standard file format mzML. To ensure the broadest possible availability, and to increase integration capabilities with popular workflow systems such as Galaxy or Nextflow, we have also built Conda and BioContainers containers around ThermoRawFileParser. In addition, we implemented a user-friendly interface (ThermoRawFileParserGUI) for those users not familiar with command-line tools. Finally, we performed a benchmark of ThermoRawFileParser and msconvert to verify that the converted mzML files contain reliable quantitative results.

## Introduction

The field of computational proteomics is approaching the big data age (1), driven both by a continuous growth in the number of samples analysed *per* experiment, as well as by the growing amount of data obtained in each analytical run. This trend towards ever more data is also directly reflected in the rapidly growing amount of publicly available proteomics data, which in turn means that there is increasing benefit to be had from the reanalysis of millions of mass spectra (2–5) to find new biological insights (e.g. novel variants and post-translational modifications (6)). However, in order to process these large amounts of (public) data, it is increasingly necessary to use elastic compute resources such as Linux-based cluster environments and cloud infrastructures (7).

The development of computational proteomics tools has historically been skewed by the development of Windows-based software tools such as ProteomeDiscover, MaxQuant (8), PeaksDB and Mascot Distiller (9). An important driver for this bias has been the lack of cross-platform libraries to access instrument output data files (RAW files) from major instrument providers (10). Several approaches have been devised to overcome this challenge, including the use of dedicated Windows machines in workflows (11) for conversion to RAW data to standard file formats such as mzML (12), the encapsulation of Windows tools such as ReAdW (13) and msconvert (14) into WineHQ (http://tools.proteomecenter.org/wiki/index.php?title=Msconvert_Wine) to make these tools Linux-compatible, and even the creation of reverse-engineered RAW file readers (15).

An important breakthrough was achieved in 2016, when Thermo Scientific released the first cross-platform application programming interface (API) that enables access to Thermo RAW files from all their instruments on all commonly used operating systems. Importantly, this provides the enticing possibility to move proteomics into Linux/UNIX environments, including scalable clusters and cloud environments. This library has already led to a new version of the popular MaxQuant framework that is compatible with Linux/UNIX environments (16), and it has also been incorporated into the cross-platform, cluster-oriented quantification tool moFF (17).

While the Thermo cross-platform library thus enables specially-developed software to access Thermo Raw files on diverse operating systems, most open-source computational proteomics workflows (e.g. OpenMS (18), Galaxy (19), and the Trans-Proteomics pipeline (TPP) (20)) are based on generic, open data formats such as Mascot Generic File (MGF) or mzML. In order to allow these tools to benefit maximally from the cross-platform access to Thermo Raw files, we here present ThermoRawFileParser, an open-source, cross-platform tool that converts Thermo RAW files into open file formats such as MGF and mzML similar to other tools such as msconvert (14) and RawTools (21). To ensure the broadest possible availability, and to increase integration capabilities with popular workflow systems such as Galaxy (22) or Nextflow (23), we have also built Conda (24) and BioContainers (25) containers around ThermoRawFileParser. Finally, we performed a benchmark of ThermoRawFileParser and msconvert to verify that the converted mzML files contain reliable quantitative results.

## Materials

### Tool Design and Integration

ThermoRawFileParser (https://github.com/compomics/ThermoRawFileParser) has been implemented following a modular design (**Figure 1**). Every file specific exporter is implemented as an independent module, which enables easy extension to include more exporters in the future. Currently, the tool can export to MGF (**MGFSpectrumWriter**), mzML (**MzMLSpectrumWriter**), and JSON (for the metadata only) (**MetadataWriter**). This modular design has already enabled the community to extend the library for other novel file formats such as Parquet (**ParquetSpectrumWriter**), which is designed for distributed big data processing clusters of Hadoop or Spark. The JSON export of ThermoRawFileParser can optionally be used to only extract various metadata elements (including instrument settings and scan settings; see https://github.com/PRIDE-Archive/pride-metadata-standard_for_the_full_list) (**Supplementary Note 1**). This specific feature is currently used by the PRIDE Database to re-annotate thousands of RAW files with the correct instrument metadata. For peak picking, data centroiding, and noise removal, ThermoRawFileParser relies on the native methods provided by the Thermo API.

**Figure 1:**
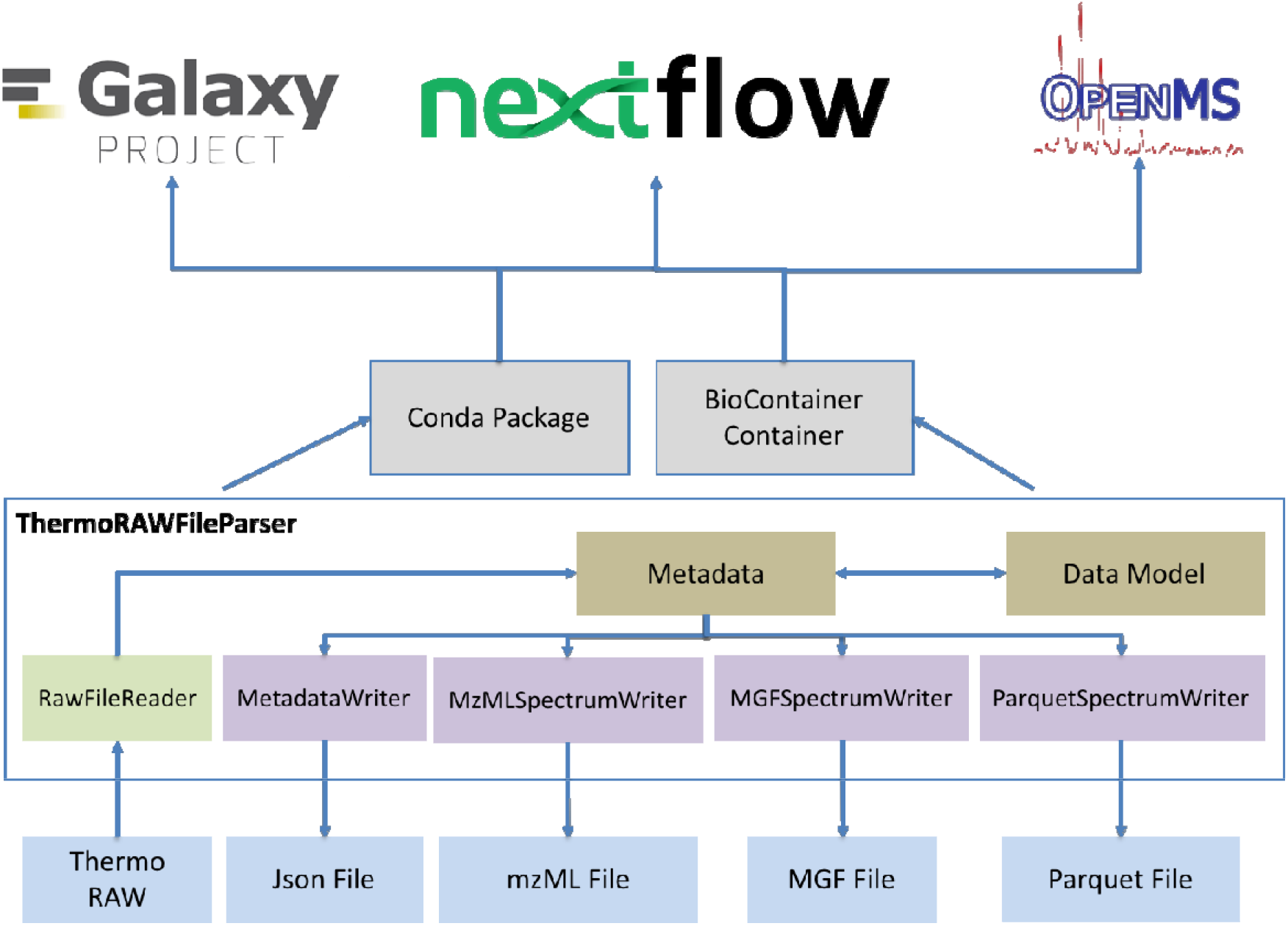
Modular design of ThermoRawFileParser includes exporters to MGF, mzML, Parquet, and Json Metadata. A Conda package and corresponding BioContainer is available for reuse in workflow engines such as Nextflow, Galaxy and OpenMS.

A key feature of any open-source tool is its ability to integrate with other frameworks (26). We have therefore created a BioConda recipe (24) for ThermoRawFileParser (https://github.com/bioconda/bioconda-recipes/tree/master/recipes/thermorawfileparser), which can be used to automatically build a Docker Container. This Docker is pushed to the BioContainer project (25), which in turn enables easy reuse of the tool by both the Galaxy (22) and the Nextflow (23) environments. As an illustration of such integration, we have developed a Nextflow workflow for the proteomics community, which converts an entire ProteomeXchange project using the ThermoRawFileParser container (https://github.com/bigbio/nf-workflows).

In addition to the command-line tool, we have implemented a graphical user interface that makes the use of ThermoRawFileParser easier and highly intuitive, enabling the user to perform conversions of RAW files. The GUI includes all main options of ThermoRawFileParser, and a report system to report errors during the conversion. ThermoRawFileParserGUI is an open source Java program, available in a cross-platform package that incorporates ThermoRawFileParser executables for the main operating systems. It can be downloaded from https://github.com/compomics/ThermoRawFileParserGUI.

### Benchmark dataset

We used the IPRG2015 dataset (https://www.ebi.ac.uk/pride/archive/projects/PXD010981) (27) to benchmark the quality of the mzML files produced by ThermoRawFileParser. This dataset is based on four artificially constructed samples of known composition, each containing a constant background of 200ng of tryptic digests of *S. cerevisiae* (ATCC strain 204508/S288c). Each sample was separately spiked in with different quantities of six individual protein digests. Samples were analysed in three LC–MS/MS using a Thermo Scientific Q-Exactive mass spectrometer (12 runs). Both MS and MS/MS data were acquired in profile mode in the Orbitrap, with resolution 70 000 for MS and 17 500 for MS/MS. The MS1 scan range was 300–1650 m/z, the normalized collision energy was set to 27%, and singly charged ions were excluded (27). The ThermoRawFileParser mzML output files are compared with mzMLs derived from msconvert (14) for benchmarking.

### Benchmark analysis workflow

To perform the benchmarking, we built a workflow using OpenMS (18, 28) in which raw files were converted from Thermo Scientific RAW files to mzML using the msconvert tool from ProteoWizard (14) on the one hand, and with ThermoRawFileParser on the other hand. The resulting spectra were centroided and searched using MS-GF+ (v2018.01.30) (29), executed via the OpenMS search engine wrapper MSGFPlusAdapter, allowing 10 ppm precursor mass tolerance, and setting carbamidomethylation of cysteine as fixed, and methionine oxidation as variable modification. All other settings were kept at their default values. PSMs were filtered (q-value < 5%) and used for feature detection using the semi-targeted approach implemented in the OpenMS tool FeatureFinderIdentification (30). Prior to identification, nonlinear retention time alignment was performed using the MapAlignerIdentification and the identified proteins were then quantified using unique peptides only (31). The workflow for comparison was developed using Nextflow (23) and BioContainers (25) to ensure the reproducibility of the present results (https://github.com/bigbio/nf-workflows/tree/master/benchmark-converter-nf).

## Results and Discussion

We compare msconvert and ThermoRawFileParser conversion to mzML using four different metrics: MS1 peak count distribution, MS2 peak count distribution, identification map, and the precursor charge distribution (**Supplementary Figure 1**). The results show major differences between both tools with regards to the number of peaks reported, and this on each MS level (MS1 and MS2). On average, the number of peaks *per* spectrum is ten-fold higher for msconvert mzML files as compared to ThermoRawFileParser mzML files. This occurs because the new peak picking method implemented in the Thermo API used by ThermoRawFileParser improved drastically with regards to the removal of noise peaks that do not contribute to identification. As a result, despite the substantial difference in the number of peaks retained, there is no major difference in the identification map and precursor charge distribution (**Supplementary Figure 1**) between the tools.

Table 1 shows the number of MS1 and MS2 spectra, and the number of identified peptides and proteins for both workflows. Across all samples and replicates, the number of identified peptides and proteins is higher for the ThermoRawFileParser workflow when compared to the msconvert workflow, despite the abovementioned higher number of peaks retained in the msconvert workflow. This identification advantage for ThermoRawFileParser derived mzML files amounts to 10% on average at the peptide level, and to 4% on average at the protein level (**Table 1**). Benchmarking protein quantification between both approaches shows no major differences between the two approaches (**Figure 2**).

**Table 1:**
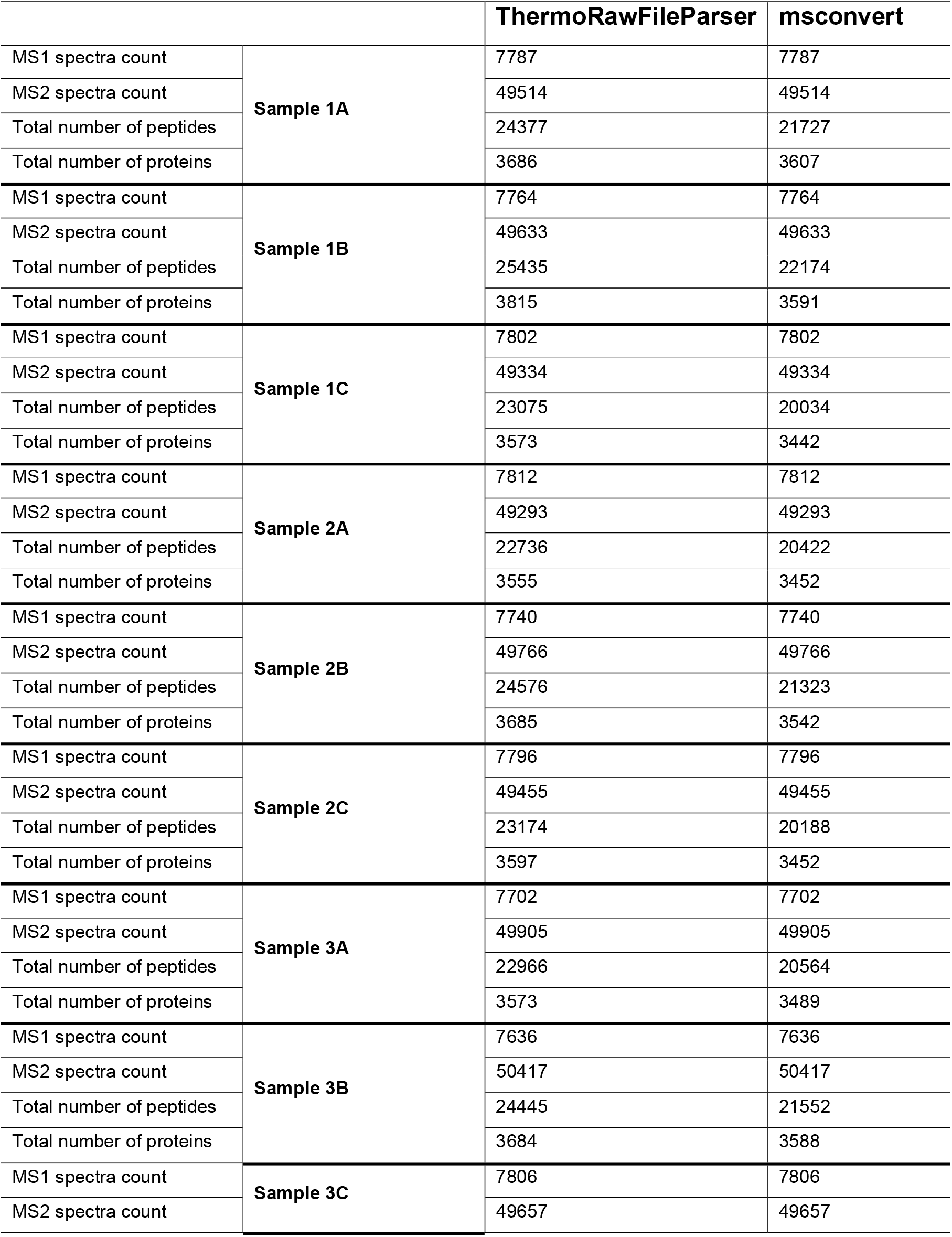
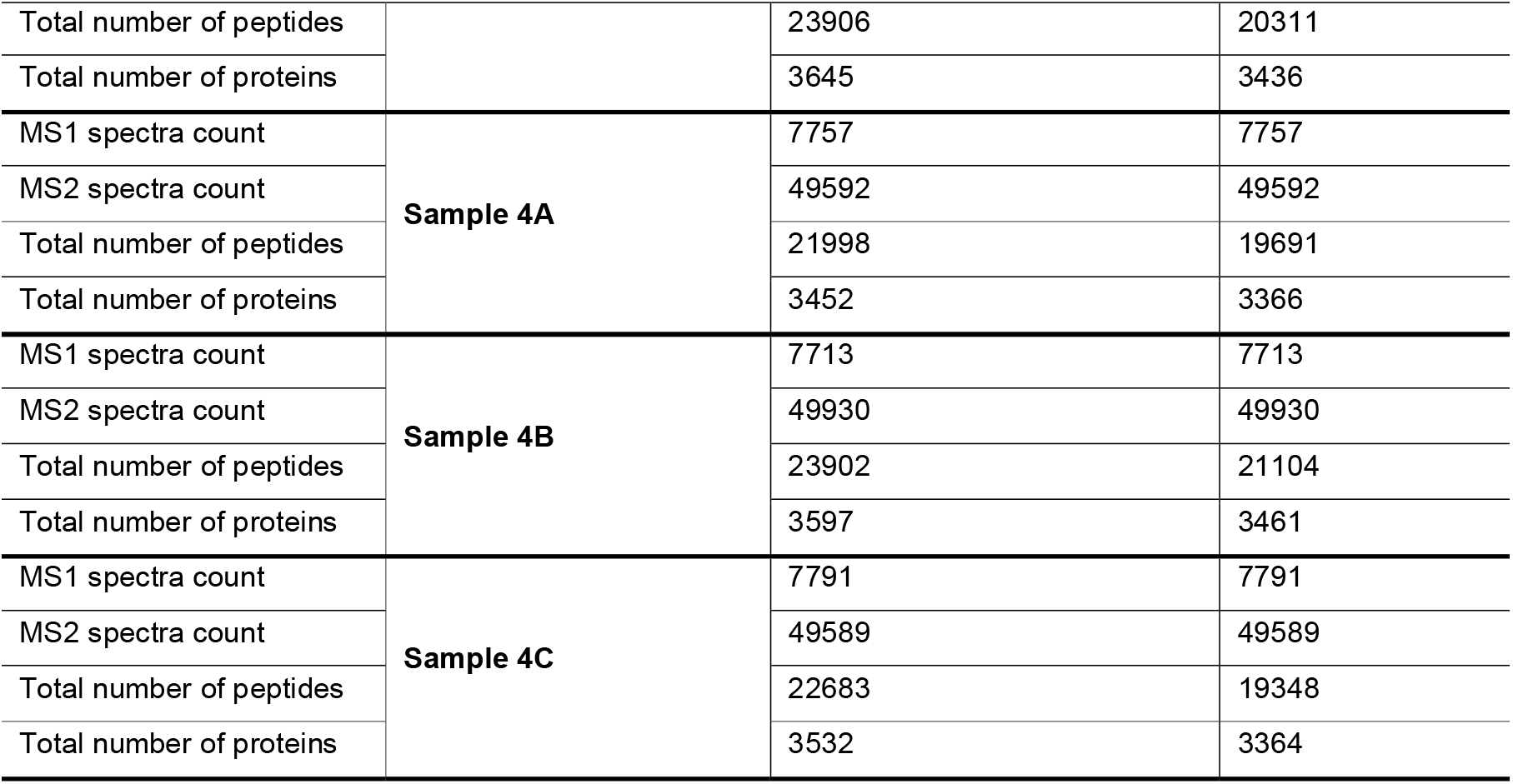
Statistics on number of spectra at MS1 and MS2 level, identified peptides, and proteins for each MS run and software workflow (ThermoRawFileParser and msconvert).

**Figure 2:**
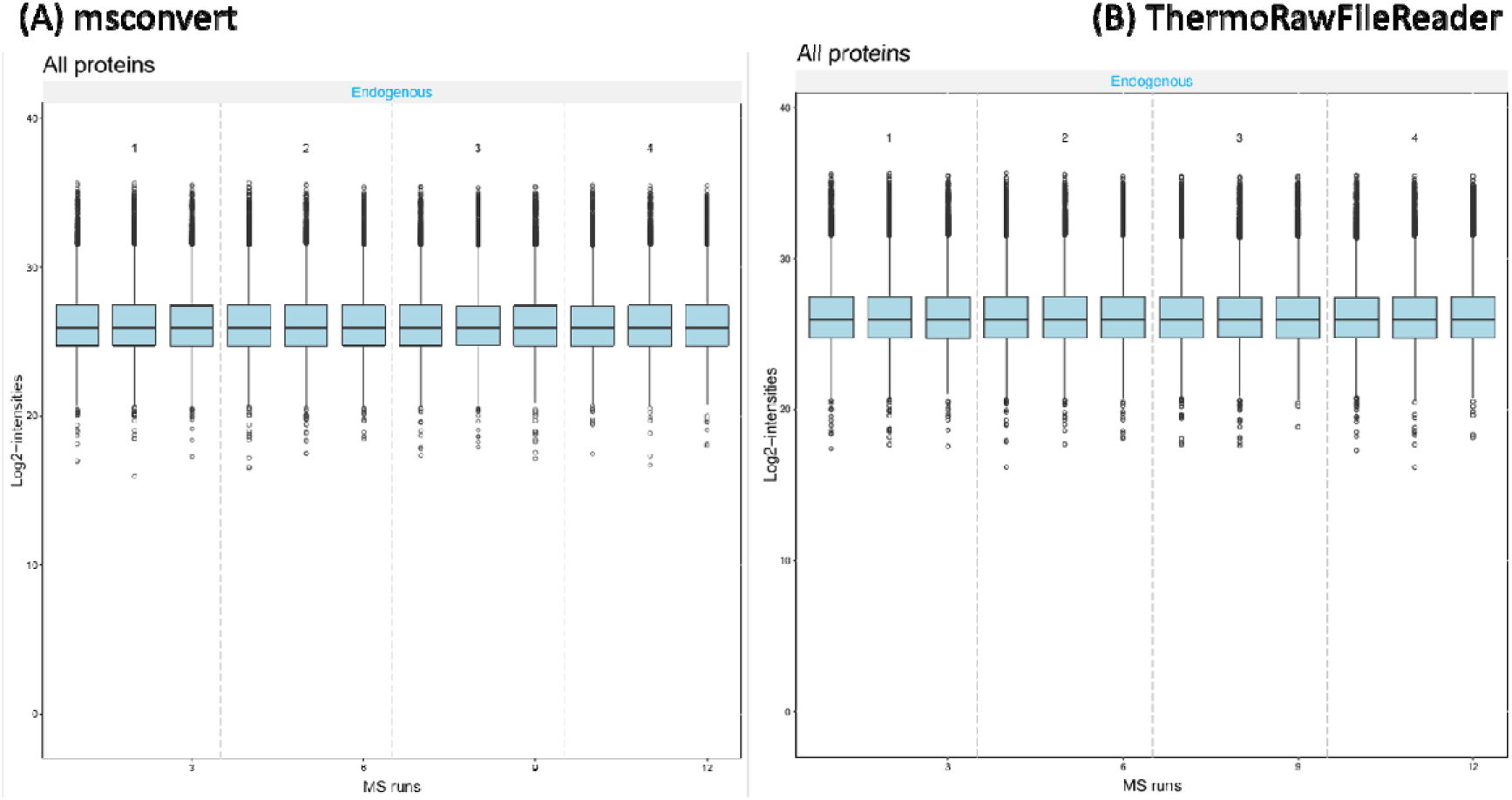
Log2-transformed intensity boxplot for four samples and twelve technical replicates for (A) msconvert-derived mzML files, and (B) ThermoRawFileParser-derived mzML files.

As a final benchmark, we analysed the IPRG2015 dataset to verify whether the mzML files obtained by the ThermoRawFileParser pipeline could replicate the quantification of the spike-in proteins in the sample using the approach described in the original publication (27). The results show that there is no appreciable difference between the IPRG 2015 analysis and the results from the ThermoRawFileParser workflow (**Figure 2**).

In addition to msconvert, the recently published RawTools (21) allows to convert RAW files into MGF files. In addition, it provides multiple options to perform QC metrics. However, RawTools is not design as a conversion tool and do not provides support for standard HUPO-PSI file formats such as mzML.

## Conclusions

ThermoRawFileParser is an open-source software tool for the conversion of Thermo Raw files into open formats. Because of the growing need for more scalable and distributed computational proteomics approaches, ThermoRawFileParser has been designed to easily plug into large-scale workflow systems such as Galaxy, Nextflow, or OpenMS. The current implementation also provides support for native writing into Amazon web service object stores (S3), making the tool highly portable to cloud architectures. Finally, the modular design of the library, along with its open source nature, allows other researchers to contribute to and extend ThermoRawFileParser for new file formats in the future. Benchmarking tests on gold standard datasets against the ProteoWizard exporter show major improvements in peak detection, and noticeable increases in peptide and protein identifications while maintaining quantitative accuracy.

## Acknowledgements

This work was partially supported by one ELIXIR Implementation Study. ELIXIR-EXCELERATE is funded by the European Commission within the Research Infrastructures programme of Horizon 2020, grant agreement numbers 676559. YP-R acknowledge the Wellcome Trust (grant number 208391/Z/17/Z) and the EPIC-XS project (grant number 823839), funded by the Horizon 2020 programme of the European Union. HB is supported by the Bergen Research Foundation and the Research Council of Norway.

