## Supplementary Information for "ThermoRawFileParser: modular, scalable and cross-platform RAW file conversion"

**Supplementary Note 1**:

**Table 1** shows the statistics about number of spectra at MS1 and MS2, identified peptides and proteins for each MS run and software workflow (ThermoRawFileParser and msconvert). **Figure 1** shows figures of the statistics about both pipelines

Table 1: QC Metrics for both converters ThermoRawFileParser and msconvert

|  | | **ThermoRawFileParser** | **msconvert** |
| --- | --- | --- | --- |
| MS1 spectra count | **Sample 1A** | 7787 | 7787 |
| MS2 spectra count |  | 49514 | 49514 |
| Total number of peptides |  | 24377 | 21727 |
| Total number of proteins |  | 3686 | 3607 |
| MS1 spectra count | **Sample 1B** | 7764 | 7764 |
| MS2 spectra count |  | 49633 | 49633 |
| Total number of peptides |  | 25435 | 22174 |
| Total number of proteins |  | 3815 | 3591 |
| MS1 spectra count | **Sample 1C** | 7802 | 7802 |
| MS2 spectra count |  | 49334 | 49334 |
| Total number of peptides |  | 23075 | 20034 |
| Total number of proteins |  | 3573 | 3442 |
| MS1 spectra count | **Sample 2A** | 7812 | 7812 |
| MS2 spectra count |  | 49293 | 49293 |
| Total number of peptides |  | 22736 | 20422 |
| Total number of proteins |  | 3555 | 3452 |
| MS1 spectra count | **Sample 2B** | 7740 | 7740 |
| MS2 spectra count |  | 49766 | 49766 |
| Total number of peptides |  | 24576 | 21323 |
| Total number of proteins |  | 3685 | 3542 |
| MS1 spectra count | **Sample 2C** | 7796 | 7796 |
| MS2 spectra count |  | 49455 | 49455 |
| Total number of peptides |  | 23174 | 20188 |
| Total number of proteins |  | 3597 | 3452 |
| MS1 spectra count | **Sample 3A** | 7702 | 7702 |
| MS2 spectra count |  | 49905 | 49905 |
| Total number of peptides |  | 22966 | 20564 |
| Total number of proteins |  | 3573 | 3489 |
| MS1 spectra count | **Sample 3B** | 7636 | 7636 |
| MS2 spectra count |  | 50417 | 50417 |
| Total number of peptides |  | 24445 | 21552 |
| Total number of proteins |  | 3684 | 3588 |
| MS1 spectra count | **Sample 3C** | 7806 | 7806 |
| MS2 spectra count |  | 49657 | 49657 |
| Total number of peptides |  | 23906 | 20311 |
| Total number of proteins |  | 3645 | 3436 |
| MS1 spectra count | **Sample 4A** | 7757 | 7757 |
| MS2 spectra count |  | 49592 | 49592 |
| Total number of peptides |  | 21998 | 19691 |
| Total number of proteins |  | 3452 | 3366 |
| MS1 spectra count | **Sample 4B** | 7713 | 7713 |
| MS2 spectra count |  | 49930 | 49930 |
| Total number of peptides |  | 23902 | 21104 |
| Total number of proteins |  | 3597 | 3461 |
| MS1 spectra count | **Sample 4C** | 7791 | 7791 |
| MS2 spectra count |  | 49589 | 49589 |
| Total number of peptides |  | 22683 | 19348 |
| Total number of proteins |  | 3532 | 3364 |

**Figure 1**: The comparison between the ThermoRawFileParser and msconvert for Sample 1,2,3,4 A fraction. Four different metrics are used MS1 peak count distribution, MS2 peak count distribution, identification map and the precursor charge distribution.

| **ThermoRawParser** | **msconvert** |
| --- | --- |
| **(MS 1 peak count)** | |
| JD_06232014_sample1_A.mzML | |
| 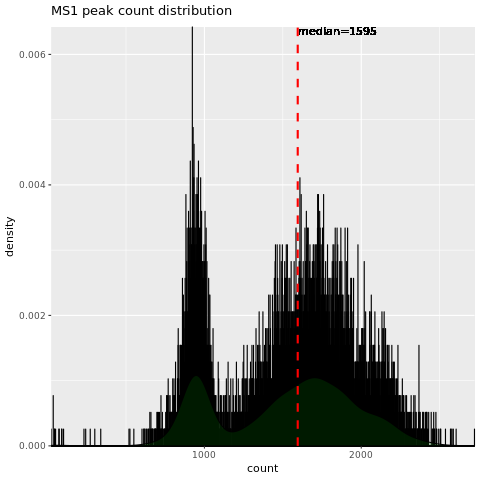 | 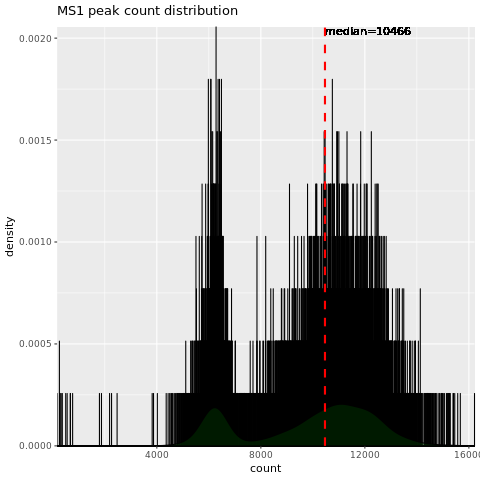 |
| JD_06232014_sample2_A.mzML | |
| 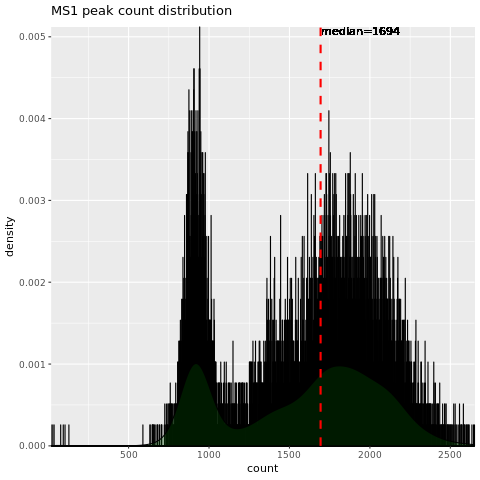 | 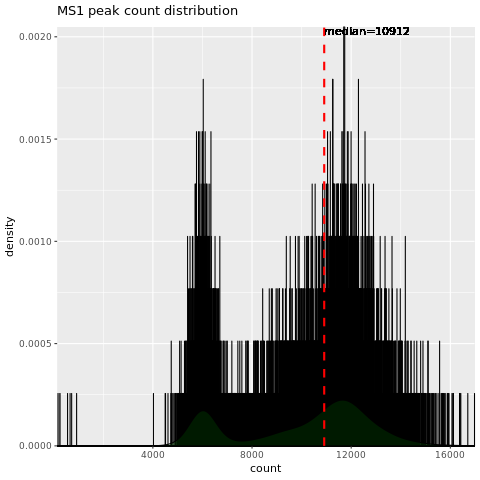 |
| JD_06232014_sample3_A.mzML | |
| 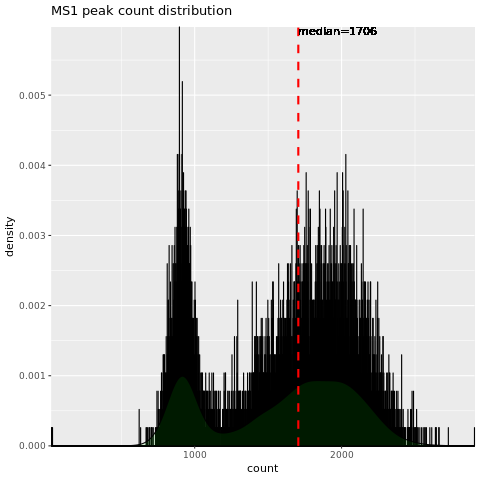 | 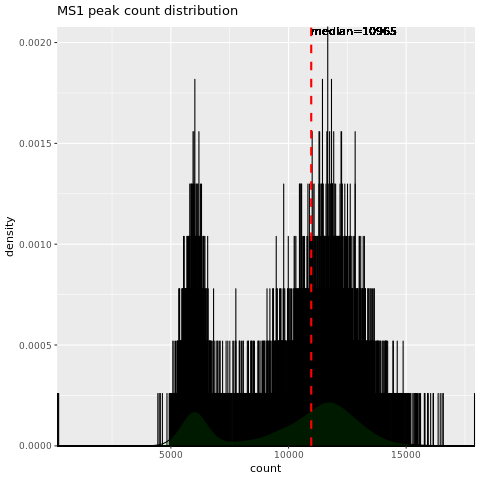 |
| JD_06232014_sample4-A.mzML | |
| 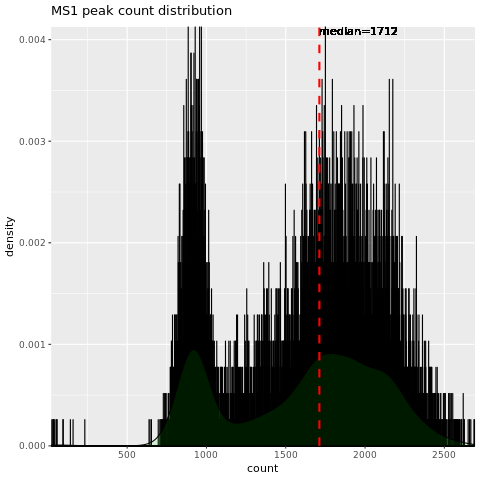 | 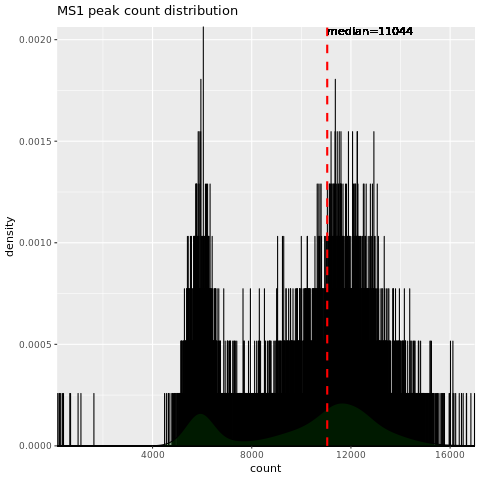 |
| **(MS 2 peak count)** | |
| JD_06232014_sample1_A.mzML | |
| 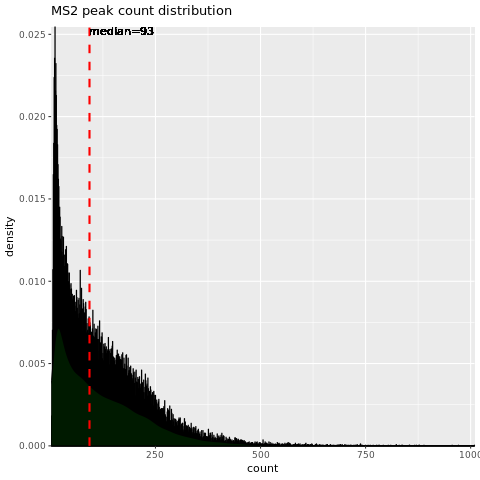 | 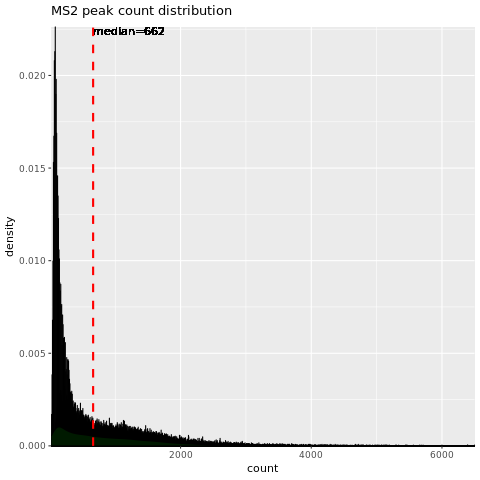 |
| JD_06232014_sample2_A.mzML | |
| 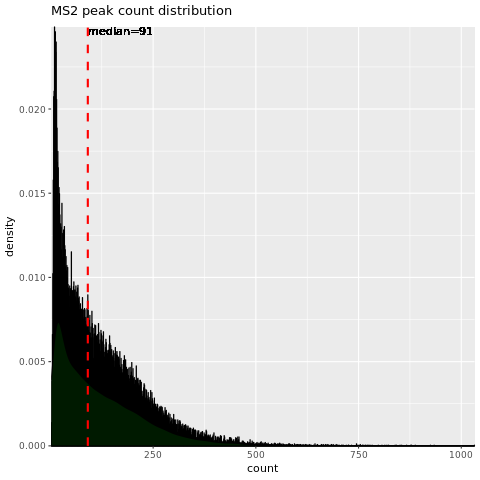 | 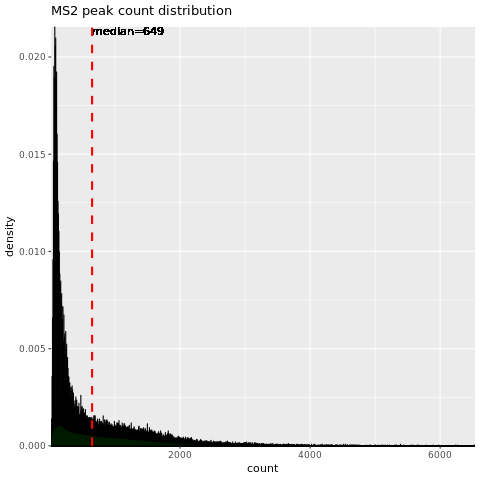 |
| JD_06232014_sample3_A.mzML | |
| 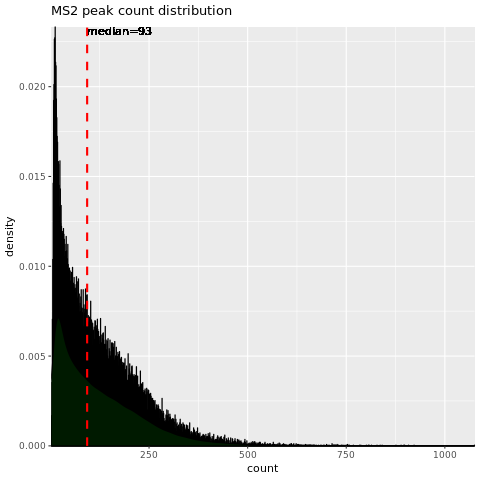 | 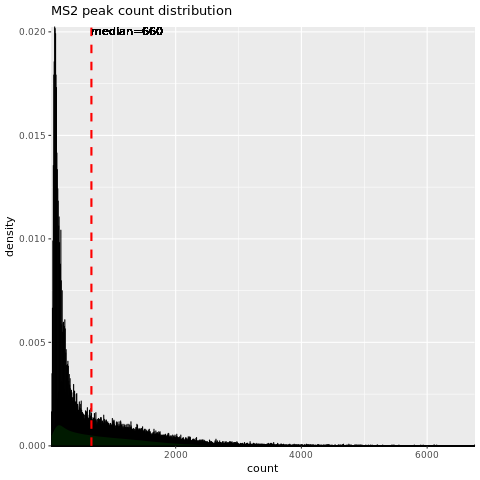 |
| JD_06232014_sample4-A.mzML | |
| 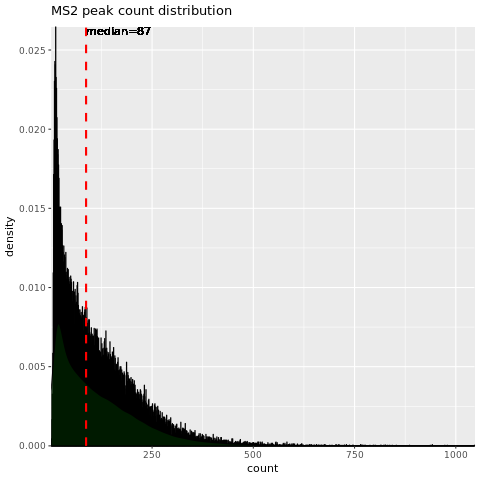 | 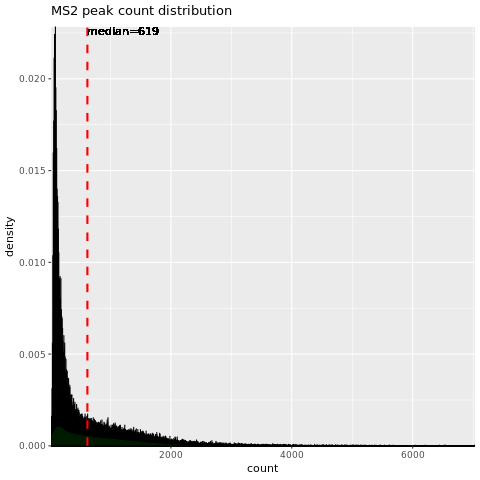 |
| **(ID Map)** | |
| JD_06232014_sample1_A.mzML | |
| 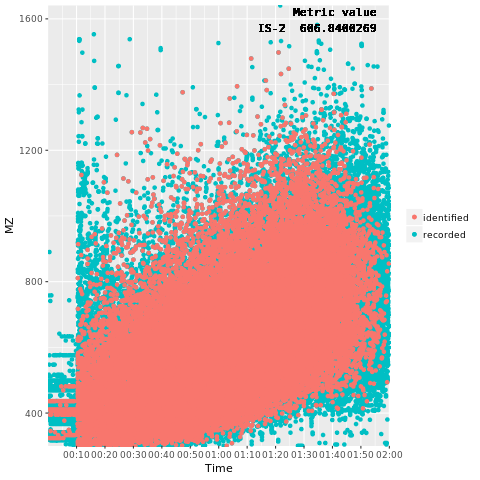 | 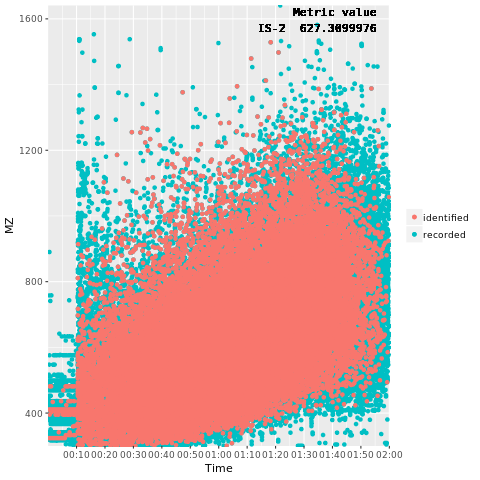 |
| JD_06232014_sample2_A.mzML | |
| 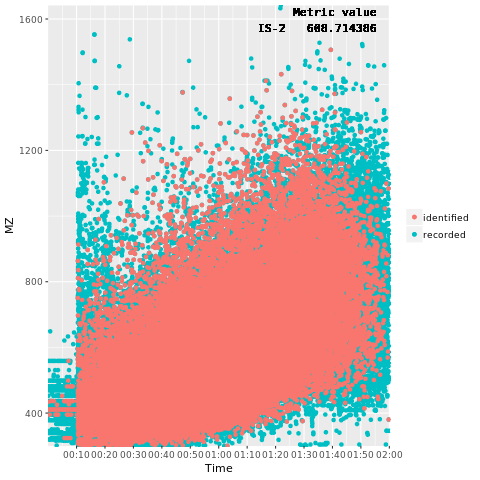 | 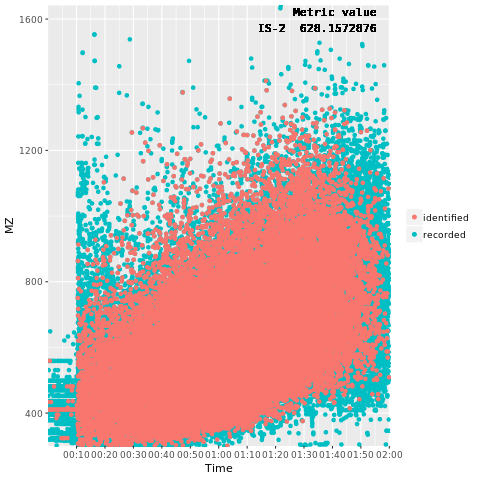 |
| JD_06232014_sample3_A.mzML | |
| 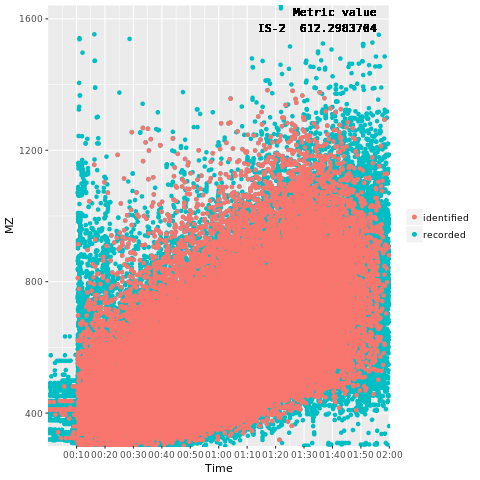 | 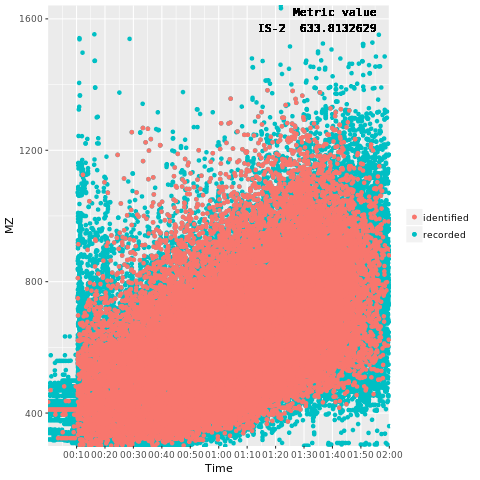 |
| JD_06232014_sample4-A.mzML | |
| 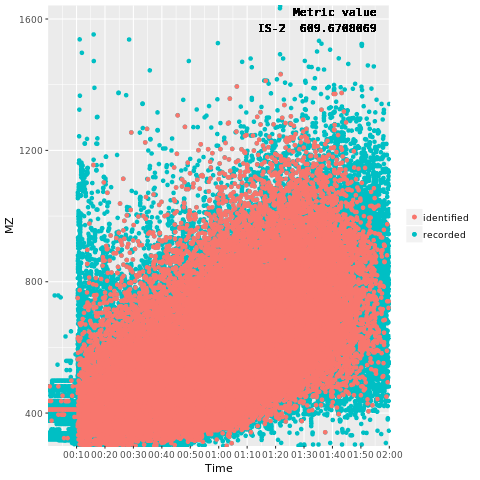 | 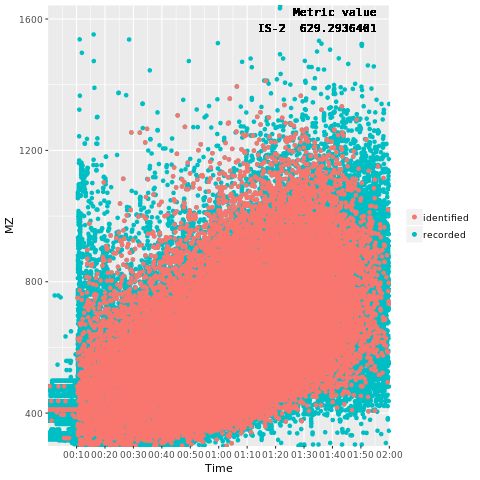 |
| **(Charge Distribution)** | |
| JD_06232014_sample1_A.mzML | |
| 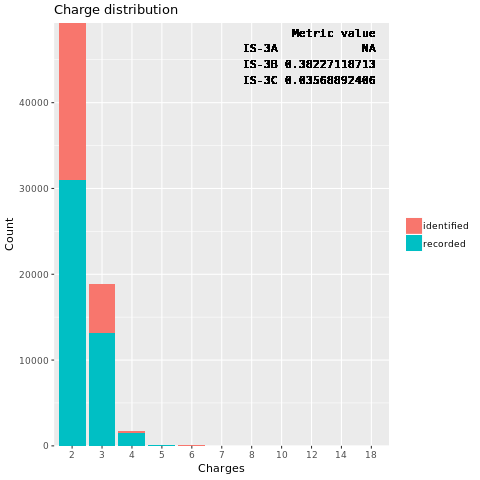 | 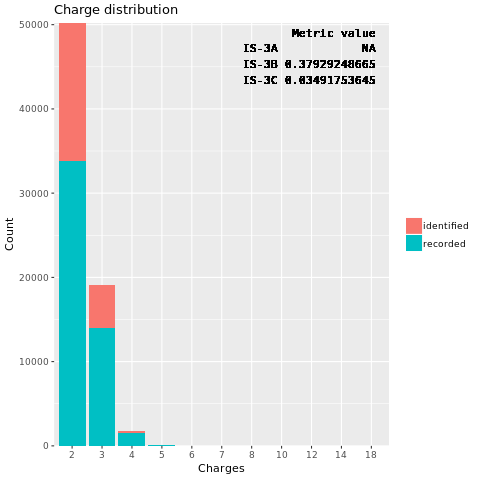 |
| JD_06232014_sample2_A.mzML | |
| 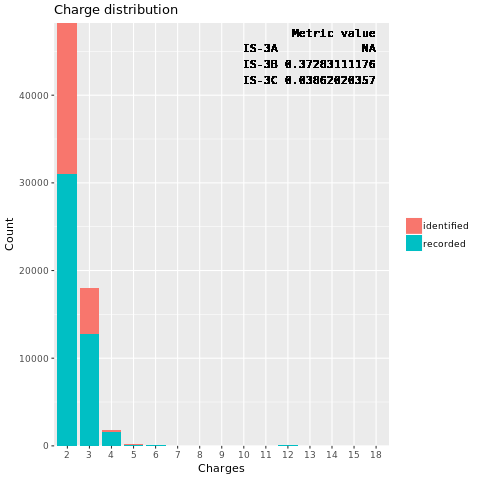 | 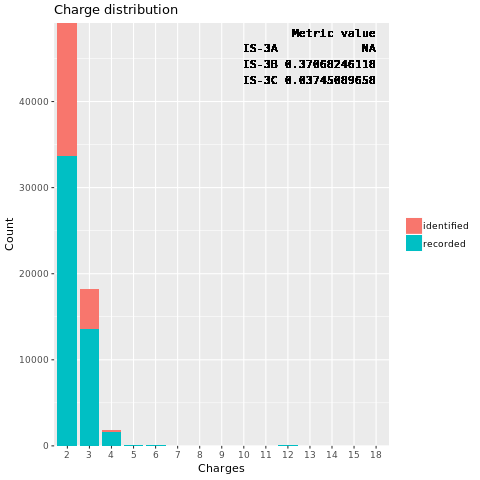 |
| JD_06232014_sample3_A.mzML | |
| 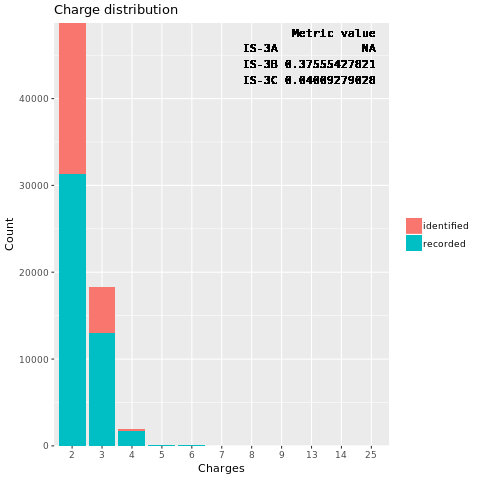 | 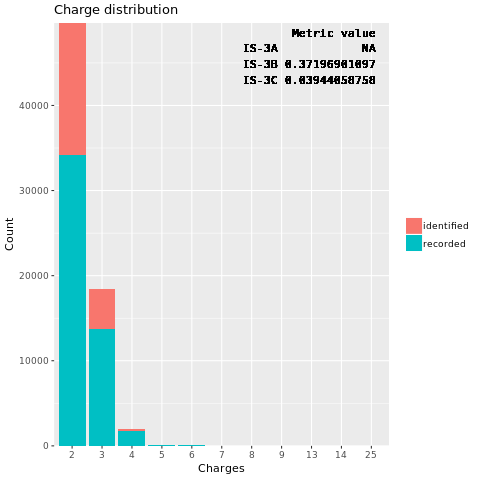 |
| JD_06232014_sample4-A.mzML | |
| 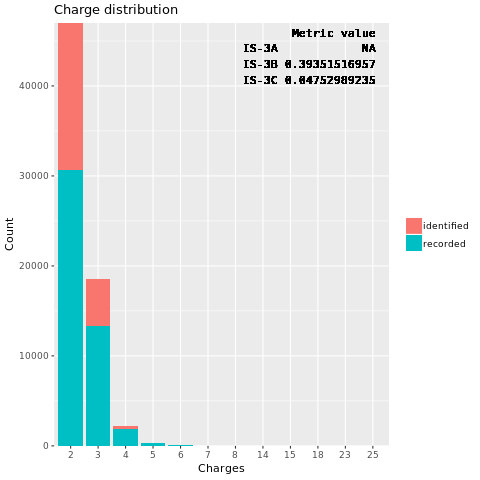 | 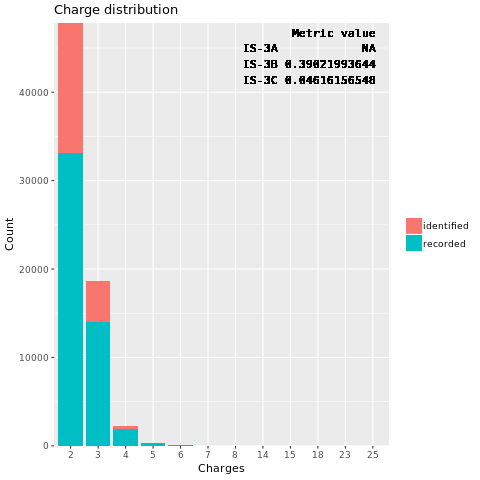 |
